# Improving the Accuracy of Forensic Age Estimation Through Bias Reduction

**DOI:** 10.64898/2026.05.30.728628

**Authors:** Maria Flores, Matteo Pellegrini

## Abstract

Chronological age estimation can provide supporting information in forensic casework when traditional identification methods are limited. DNA methylation, a stable epigenetic mark, has emerged as a promising tool for predicting chronological age from trace samples. However, many existing age estimation models rely on linear regression approaches, which often yield biased prediction errors across the age distribution (i.e. model residuals show a significant age dependence). In this study, we compared three approaches for age estimation modeling: multivariable linear regression, random forest regression and maximum likelihood estimation. While the first two approaches are well established, for the third one we constructed and validated a DNA methylation-based LOESS regression maximum likelihood model for age estimation utilizing forensic-relevant CpG markers. In all cases, model performance was evaluated through Leave-One-Out Cross-Validation (LOOCV). We utilized three independent publicly accessible methylation datasets collected using droplet digital PCR (ddPCR) to evaluate the most effective method for accuracy and bias in age estimation. Notably, when we compare the results of the maximum likelihood approach to the other approaches, multivariable linear regression and random forest regression, we find less bias in the age associated residuals compared to the other methods. These findings highlight the utility of non-linear modeling techniques in reducing the biases of epigenetic age estimation for forensic applications.

## Introduction

DNA methylation patterns used for the construction of epigenetic clocks are a promising avenue for age prediction of individuals in judicial, civil, and criminal investigations. An epigenetic age estimator is a mathematical model developed through machine learning techniques that regresses chronological age against a curated subset of CpG sites demonstrating the highest predictive value for age estimation. These models quantify epigenetic age by translating the methylation patterns of the selected CpG sites into temporal units expressed in years [1]. Numerous studies have developed distinct methodologies for age prediction across multiple tissues, with the most prominent epigenetic clocks being those established by Horvath [2], Hannum [3], DNAm PhenoAge [4], and DNAm GrimAge [5].

DNA methylation occurs through the covalent attachment of a methyl group to the 5’ carbon position of cytosine residues throughout the genome [6]. Most techniques involved in DNA methylation analysis use DNA conversion with sodium bisulfite allowing a chemical modification of non-methylated cytosines (C) to uracils, while the methylated cytosines remain intact [7]. This technique has been evaluated for age prediction in previous studies [8-11]. Converted cytosines can be detected by either sequencing, microarrays or droplet digital PCR (ddPCR). ddPCR has been employed in several studies that sought to develop age estimators for forensic applications because it can be applied to samples with low amounts of DNA [12-14]. ddPCR involves partitioning a sample into thousands of micro-droplets, each containing single or few copies of the target molecules, followed by amplification via PCR. Given that PCR amplification occurs concurrently within numerous micro-droplets, this technique mitigates PCR bias between methylated and non-methylated strands commonly observed in conventional PCR. The high precision of ddPCR in quantifying DNA methylation renders it a straightforward and practical method for forensic laboratories to analyze small numbers of CpG markers [12].

Additionally, most DNA methylation–based age prediction models, or “epigenetic clocks,” are built using linear regressions of methylation levels at selected CpG sites against chronological age. However, methylation changes at individual CpG sites are often nonlinear across the lifespan, with rapid alterations during development followed by slower or plateauing changes in adulthood and late life [2-3, 19]. When this nonlinearity is not properly modeled, systematic biases arise in the residuals of age predictions: young individuals tend to be underpredicted, middle-aged individuals overpredicted, and older individuals again underpredicted [18]. These residual patterns reflect the inadequacy of purely linear models to capture the exponential or piecewise nature of methylation trajectories at many CpGs. More recent approaches, such as the Epigenetic Pacemaker model [19], explicitly account for nonlinear methylation dynamics by fitting site-specific rates of change, thereby reducing these age-dependent residual biases and better capturing the biological tempo of epigenetic aging. Another approach we developed previously for bisulfite sequencing data called BayesAge explicitly models the nonlinear trends of each individual CpG site with age and uses maximum likelihood to estimate age[17].

In this study, we seek to compare different approaches for estimating age using DNA methylation data: multiple/multivariable regression, random forest regression and a maximum likelihood approach. For the latter method, we present a non-linear age estimation framework based on DNA methylation using CpG markers relevant to forensic applications. Methylation data obtained from ddPCR were examined across three independent datasets: the first evaluated markers ELOVL2, PDE4C, and FHL2 [12]; the second evaluated ASPA, IGSF11, MEIS1-AS3, and COL1A1 [14], and the third evaluated ELOVL2, TRIM59, and FHL2 [15]. We show that the maximum likelihood approach generates age estimates with reduced age associated biases compared to the other two approaches, thus presenting a promising alternative for forensic age estimation.

## Methods

### Data Acquisition

The first dataset examined in this study was obtained from Dias and Manco et al. [12] and consisted of blood-derived DNA methylation (DNAm) samples from 58 Portuguese individuals, including 42 females and 16 males. The participants’ ages ranged from 1 to 93 years, with a mean age of 41.91 years. Specifically, 12 individuals were aged between 1 and 19 years, 15 individuals between 20 and 39 years, 17 individuals between 40 and 59 years, and 14 individuals between 60 and 93 years. The second dataset analyzed was obtained from Zhou et al. [14] and consisted of blood-derived DNAm samples from 327 unrelated individuals residing in Chengdu, Sichuan Province, China, with ages spanning from 20 to 100 years. The third dataset was obtained from Fakhr et al. [15] and consisted of 250 blood samples obtained from the National Biobank of Korea with ages spanning from 20 to 69 years old.

### Statistical Analysis

All data analysis was conducted using RStudio (version 4.3.2) and the residuals of all models were examined to assess their age-dependent biases. Leave-One-Out-Cross-Validation (LOOCV) was utilized as the primary method to evaluate the performance of all models. In this specific approach, a single individual sample was excluded from the entire dataset, and the model was then trained exclusively on the remaining samples, with the excluded sample being used as the test instance to assess predictive accuracy. This procedure was repeated iteratively for every sample in the dataset to ensure a comprehensive evaluation of model performance (Fig. 1). All models were evaluated using mean absolute error (MAE) as the performance metric. For a quantitative comparison of the residuals, the absolute value of the area beneath the residual curves was quantified.

**Figure 1.**
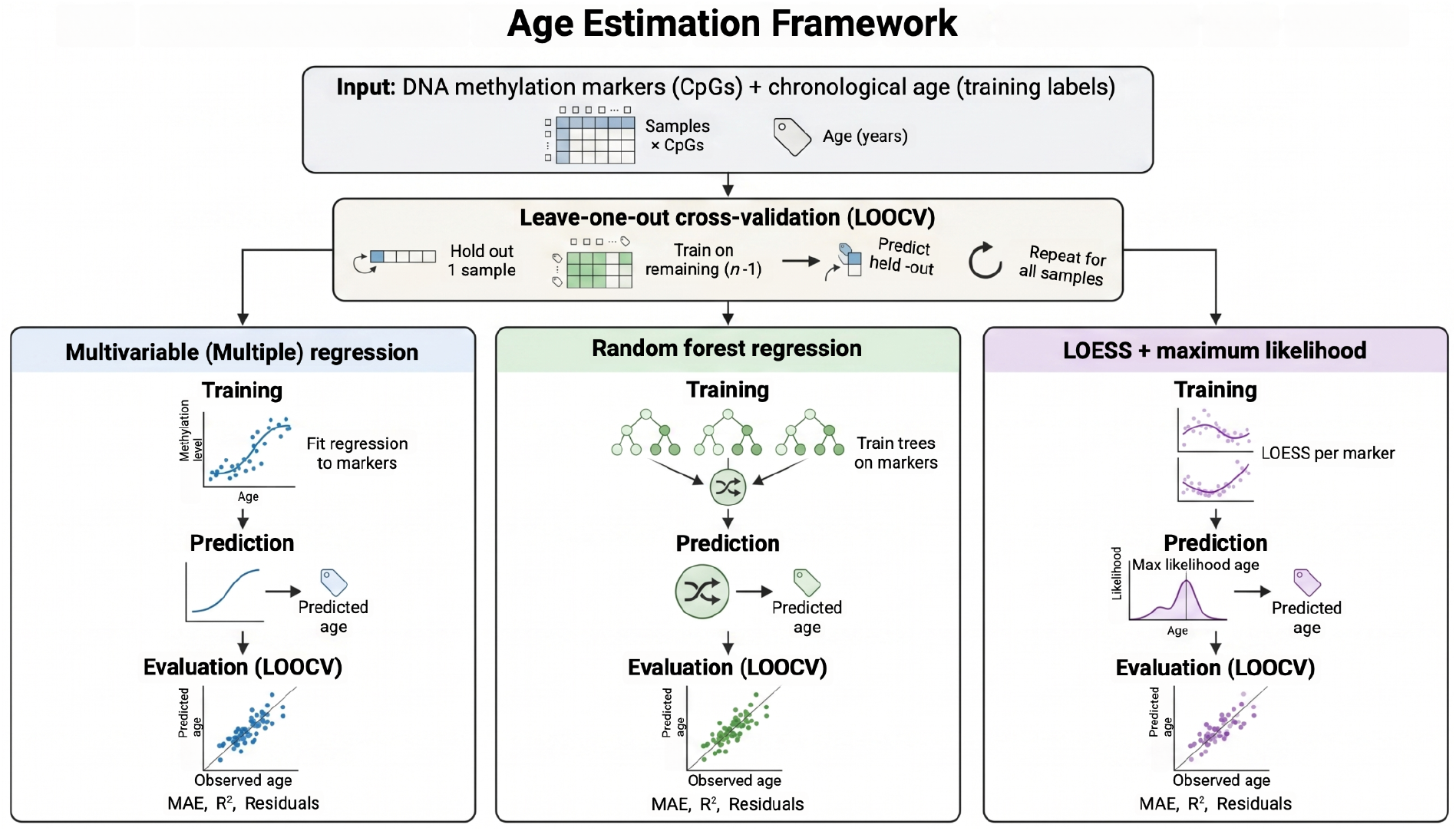
Age prediction overview

### Multiple/Multivariable Regression Analysis

The dataset from Dias and Manco [12] utilized a multivariable linear regression model incorporating DNAm data from three markers: ELOVL2 (modeled as a quadratic term), PDE4C, and FHL2. The age prediction equation is defined as: −25.166 + 0.003 × (ELOVL2 methylation)^2^ + 0.924 × (FHL2 methylation) + 0.527 × (PDE4C methylation). In contrast, the dataset from Fakhr et al. [15] employed a multiple regression analysis involving three markers: ELOVL2, FHL2, and TRIM59. Their age prediction formula is expressed as: intercept + (coefficient × ELOVL2) + (coefficient × FHL2) + (coefficient × TRIM59). Given that the original publications did not disclose their source code, the published models were reproduced by refitting the same regression framework. Multiple/multivariable regression was implemented using the *lm* function in R.

### Random Forest Regression Analysis

The dataset from Zhou et al. [14] employed a random forest regression model in Python utilizing DNA methylation data from four markers: ASPA, IGSF11, COL1A1, and MEIS1-AS3. The authors indicated that the same code as Jiang et al. [20] was used; however, the source code from the original study was also not made publicly available. The only detail provided was that the analysis was conducted using 20 decision trees. Here, random forest regression was implemented using the *ranger* function in R using 20 decision trees.

### Maximum Likelihood Analysis

LOESS regression was implemented using the *LOESS* function in R, as a flexible and robust method to model the non-linear relationship between DNA methylation levels measured for each marker and the chronological age of the samples. The degree of smoothness and flexibility of the fitted regression curve is primarily controlled by the *span* parameter. This parameter dictates the proportion of data points used to fit each local regression, with smaller values leading to less smoothing and larger values resulting in a smoother fit [16]. A predetermined span value of 0.75 was implemented in this model to effectively balance the trade-off between overfitting, which can occur with smaller span values, and underfitting, which is more common with span values approaching one.

The procedure for predicting age involves calculating the likelihood of observing the specific methylation levels at a given age *A* across all DNA methylation markers (as seen in (1)) by computing the product of the individual probabilities associated with each marker’s methylation level at that age (as seen in (2)). To maintain numerical stability and avoid computational issues such as underflow, this product is converted into a sum of log-likelihoods (as seen in (3)). Finally, the predicted age for a given sample is obtained by identifying the age value that maximizes this cumulative sum of log-likelihoods across all markers which can then be interpreted as the most probable estimate of the sample’s chronological age (as seen in (4)).

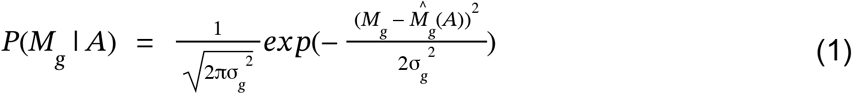

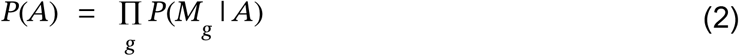

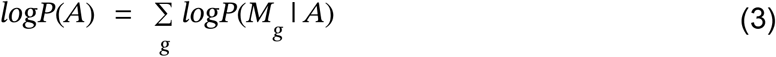

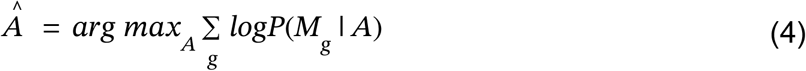

Here, *M*_g_ is the observed methylation value for gene 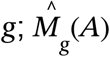 is the predicted methylation at age 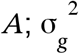 is the variance for gene 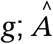 is the estimated age maximizing the likelihood.

## Results

We aimed to compare various methods for estimating age using DNA methylation data: multiple/multivariable regression, random forest regression, and a maximum likelihood approach. For the latter technique, we introduced a non-linear age estimation model based on DNA methylation utilizing CpG markers pertinent to forensic applications. Methylation data obtained from ddPCR were analyzed across three separate datasets: the first assessed markers ELOVL2, PDE4C, and FHL2 [12]; the second assessed ASPA, IGSF11, MEIS1-AS3, and COL1A1 [14]; and the third assessed ELOVL2, TRIM59, and FHL2 [15].

In the Dias et al. [12] dataset, ELOVL2 demonstrated a non-linear increase with age, approaching saturation in older age groups. PDE4C exhibited a more moderate linear association with age, accompanied by greater variability, while FHL2 showed a distinct positive linear trend with age and a relatively consistent dispersion (Fig. 2). We constructed three distinct models to predict the age of each individual from the methylation of the three markers. The multivariable model achieved a mean absolute error (MAE) of 5.14 years and an R^2^ of 0.929. The random forest model yielded comparable results with a slightly larger MAE of 5.56 years and an R^2^ of 0.906. The maximum likelihood model outperformed both, attaining an MAE of 3.73 years and an R^2^ of 0.957 overall (Fig. 2). Analysis of residual patterns revealed that the multivariable model exhibited a non-linear trend indicative of bias across age ranges, characterized by underestimation and overestimation. The random forest model mitigated this bias relative to the multivariable approach but still displayed curvature, particularly in older age groups. Conversely, the maximum likelihood model presented residuals that were more symmetrically distributed around zero, reflecting minimal bias despite a slight increase in error at advanced ages (Fig. 2).

**Figure 2.**
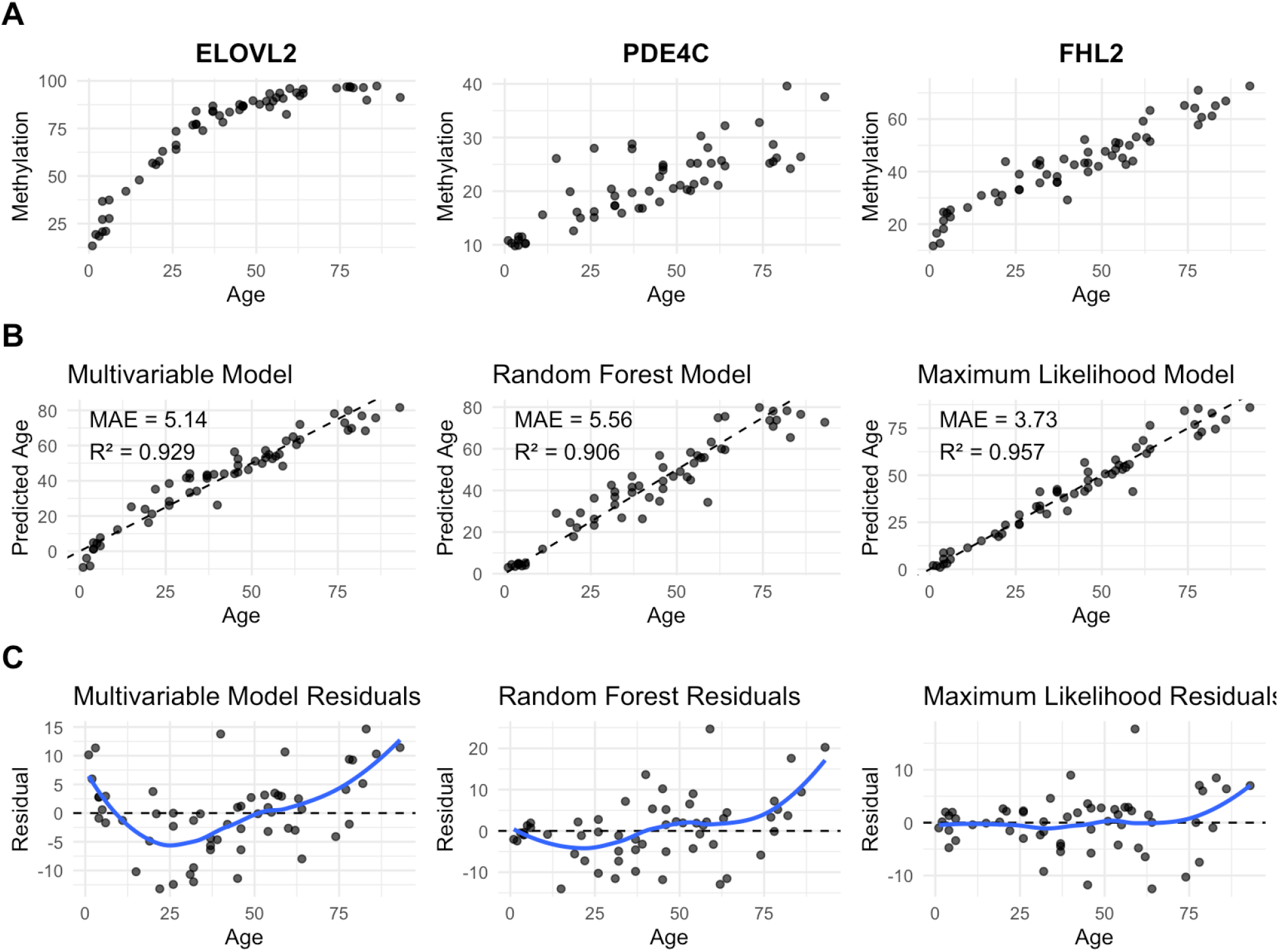
Comparison of multivariable regression, random forest regression, and maximum likelihood approach. (A) Scatterplots showing relationship between chronological age and methylation levels for markers ELOVL2, PDE4C, and FHL2. (B) Predicted versus observed age plots generated from multivariable regression, random forest regression, and the LOESS-based maximum likelihood framework. (C) Residual plots for each model showing prediction error across chronological age.

In the Zhou et al. [14] dataset, markers ASPA, IGSF11, and MEIS1-AS3 display a negative relationship with age with methylation decreasing as age increases (Fig. 3). To optimize the visualization of the plot, the fourth marker scatterplot (COL1A1) is provided in the supplementary materials (Fig. S1). Regarding model performance, the multivariable model achieved a MAE of 5.47 years and an R^2^ of 0.830. The random forest model achieved a slightly improved MAE of 5.05 years and an R^2^ of 0.853. The maximum likelihood model performed the worst attaining an MAE of 6.64 years and an R^2^ of 0.742 (Fig. 3). Analysis of residual patterns revealed that the multivariable model underestimates ages in younger age groups while overestimating them in older ages. The random forest model reduced this bias relative to the multivariable approach, although it still exhibited some curvature, particularly among older age groups. Although the maximum likelihood model demonstrated the highest overall prediction error, it showed improved bias performance compared to the other models, with only a slight increase observed in the older age groups (Fig. 3).

**Figure 3.**
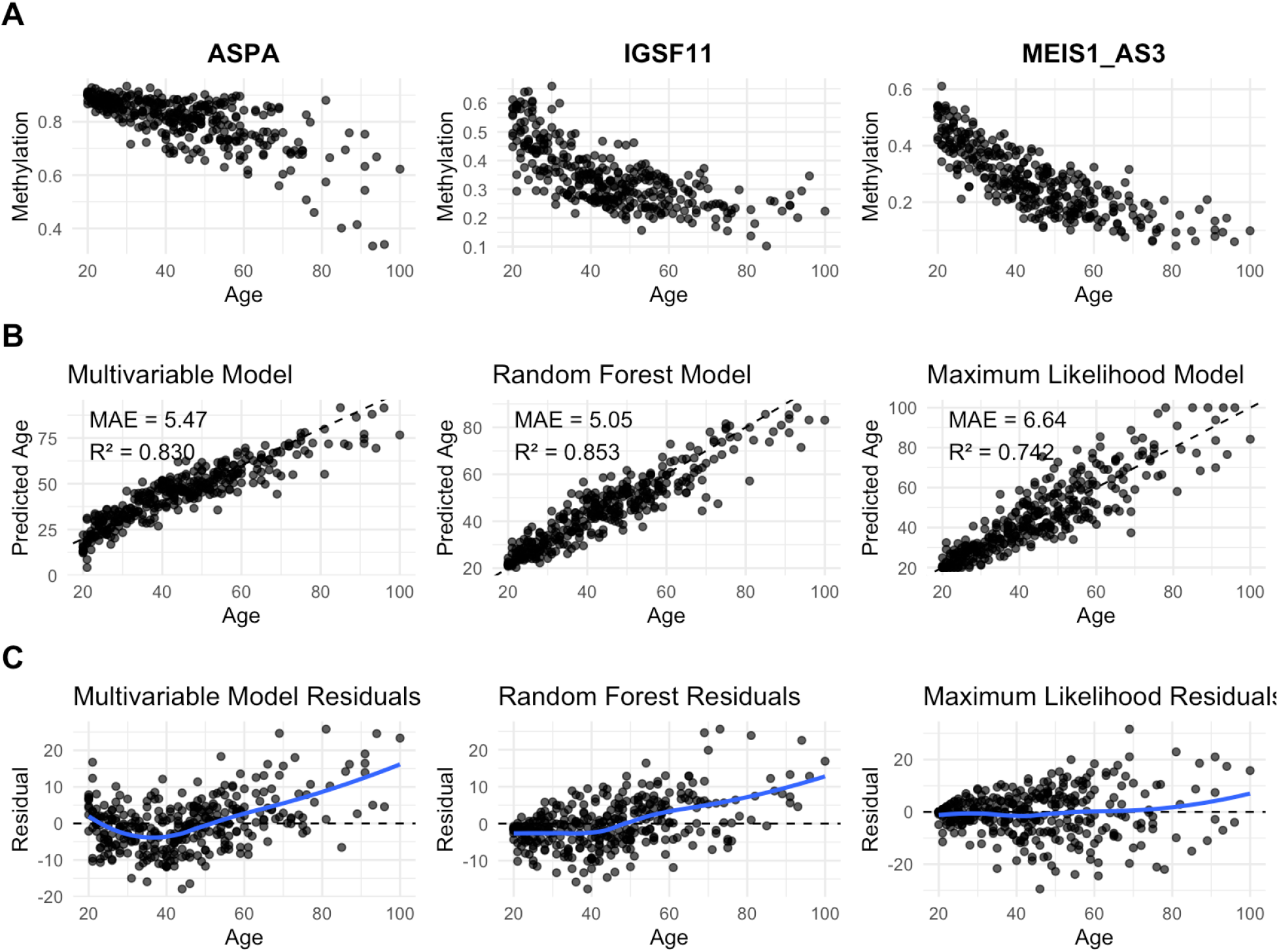
Comparison of multivariable regression, random forest regression, and maximum likelihood approach. (A) Scatterplots showing relationship between chronological age and methylation levels for markers ASPA, IGSF11, and MEIS1-AS3 (see Figure S1 for COL1A1 scatterplot). (B) Predicted versus observed age plots generated from multivariable regression, random forest regression, and the LOESS-based maximum likelihood framework. (C) Residual plots for each model showing prediction error across chronological age.

In the Fakhr et al. [15] dataset, markers ELOVL2, FHL2, and TRIM59 display a positive relationship with age with methylation increasing as age increases (Fig. 4). Regarding model performance, the multivariable model achieved a MAE of 3.09 years and an R^2^ of 0.919. The random forest model achieved a slightly worse MAE of 3.27 years and an R^2^ of 0.904. The maximum likelihood model attained an MAE of 3.04 years and an R^2^ of 0.915 (Fig. 4). Analysis of residual patterns show that the multivariable model displays a minor underestimation in younger ages and an overestimation in older ages. Similarly, the random forest model displays a minor underestimation in younger ages and a slight overestimation in older ages. The maximum likelihood model presented residuals that were more symmetrically distributed around zero, reflecting minimal bias (Fig. 4).

**Figure 4.**
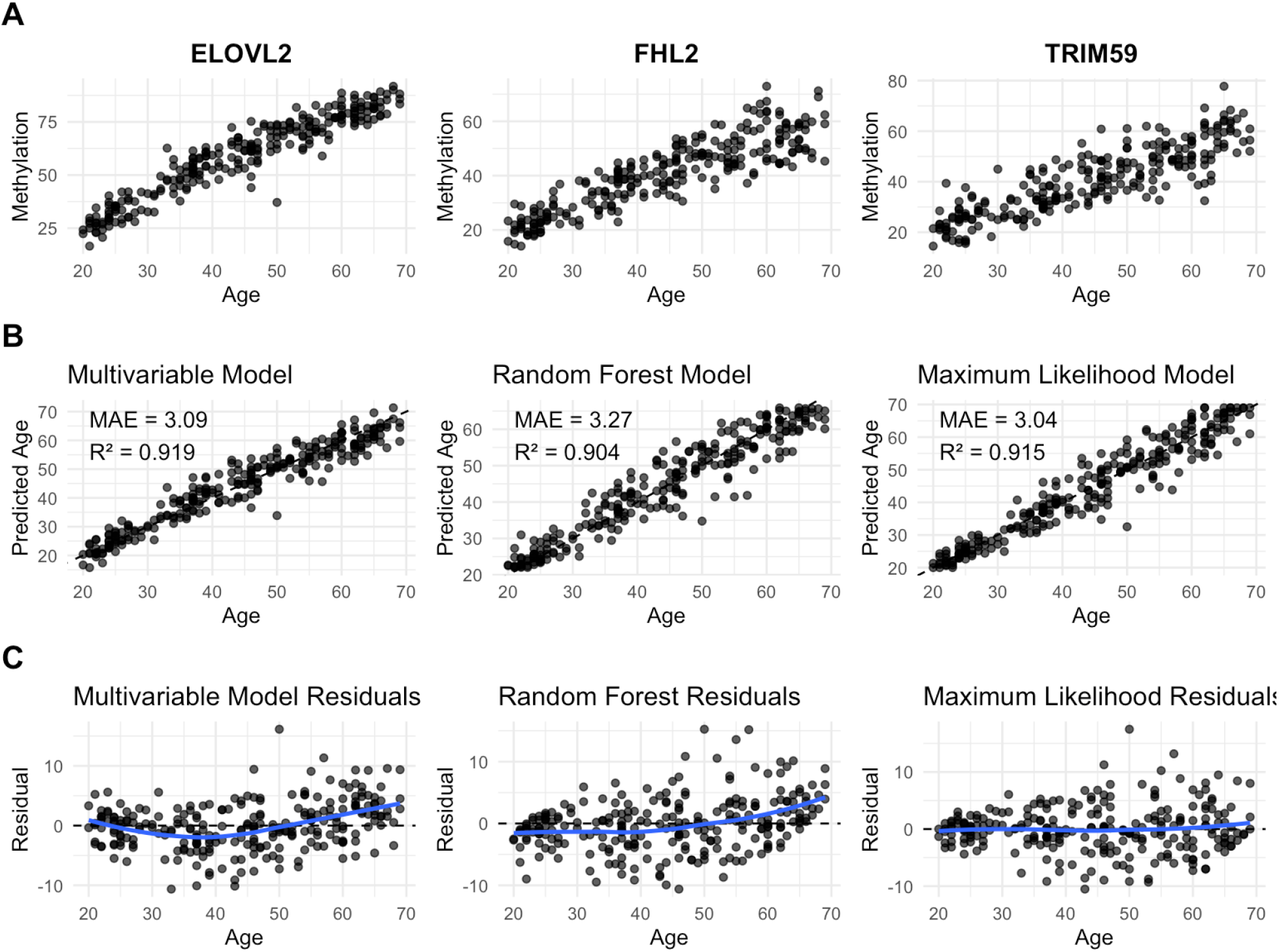
Comparison of multivariable regression, random forest regression, and maximum likelihood approach. (A) Scatterplots showing relationship between chronological age and methylation levels for markers ELOVL2, FHL2, and TRIM59. (B) Predicted versus observed age plots generated from multivariable regression, random forest regression, and the LOESS-based maximum likelihood framework. (C) Residual plots for each model showing prediction error across chronological age.

To quantitatively assess the residuals, the absolute value of the area under the residual curves was calculated and is presented in Table 1. Across all datasets, the LOESS combined with the Maximum Likelihood framework yielded the lowest residual curve area values, indicating the minimal residual bias across age ranges relative to the other modeling techniques. Specifically, for the Dias & Manco dataset [12], the LOESS + Maximum Likelihood model markedly outperformed both multivariable regression and random forest regression, with an absolute area value of 90.80343 compared to 355.0049 and 341.8449, respectively. Similarly, in the Zhou et al. dataset [14], the LOESS + Maximum Likelihood approach exhibited the lowest residual curve area at 128.3835, outperforming multivariable regression and random forest regression, which had values of 442.1757 and 384.7516, respectively. For the Fakhr et al. dataset [15], all models produced comparatively lower residual curve areas than those observed in the other datasets, with the LOESS + Maximum Likelihood model again achieving the lowest value at 10.77696.

**Table 1.** Normalized absolute area under the residual curve across datasets and age prediction models.

| **Dataset** | **Multivariable**<br>**(Multiple)**<br>**Regression** | **Random Forest**<br>**Regression** | **LOESS +**<br>**Maximum**<br>**Likelihood** |
| --- | --- | --- | --- |
| Dias & Manco | 355.0049 | 341.8449 | 90.80343 |
| Zhou et al. | 442.1757 | 384.7516 | 128.3835 |
| Fakhr et al. | 69.52811 | 67.67542 | 10.77696 |

## Discussion

ddPCR has been applied in various studies aimed at creating age prediction tools for forensic purposes due to its suitability for samples with low DNA input [12-14]. The high accuracy of ddPCR in measuring DNA methylation makes it a simple and practical technique for forensic laboratories to evaluate a small number of CpG markers [12]. Furthermore, most DNA methylation–based age prediction models are constructed using linear regression of methylation levels at selected CpG sites against chronological age. However, methylation changes at individual CpG sites frequently exhibit nonlinear patterns throughout life, featuring rapid shifts during development followed by slower or stabilized changes in adulthood and later years [2-3, 19]. Failure to properly account for this nonlinearity leads to systematic errors in age prediction residuals: younger individuals are typically underpredicted, middle-aged individuals overpredicted, and older individuals underpredicted once more [18].

In this study, we evaluated a comparison of different approaches for estimating age using published ddPCR DNA methylation data: multiple/multivariable regression, random forest regression, and a maximum likelihood method. The latter methodology employed LOESS smoothing to effectively model the non-linear association between age and methylation levels, alongside a maximum likelihood framework for age prediction.

The model consistently demonstrated reduced residual age associated bias relative to the alternative approaches. The absence of residual bias signifies that the model’s predictions do not systematically overestimate or underestimate the true values across the data. This is critical, as such bias would result in skewed age predictions. Consequently, this suggests that the model’s predictions are generally accurate and do not exhibit a tendency to favor either higher or lower values. The residual analyses presented here demonstrated that model performance should not be evaluated solely, as systematic age-dependant prediction bias remained evident in some models despite strong overall accuracy metrics.

The evaluation of both prediction accuracy and residual bias is critical in the development and assessment of age estimation models. As model performance advances, their practical implementation in real-world forensic applications becomes increasingly feasible. For instance, the VISAGE project, conducted in collaboration with Erasmus Medical Center and Polish researchers from Jagiellonian University and the Central Forensic Laboratory of the Police in Warsaw, reports the creation of a highly sophisticated forensic DNA methylation tool capable of estimating chronological age from biological samples with an accuracy typically within approximately three years. The tool has reportedly undergone rigorous validation and is currently being tested internationally across diverse populations prior to routine forensic deployment. More recently, a subsequent European initiative, termed forMAT, was launched in September 2025 to further enhance forensic methylation analysis through improved age prediction accuracy, the identification of minors in migration contexts, and the integration of lifestyle and behavioral profiling methodologies [21].

The article [21] further emphasizes the ELOVL2 marker as a robust and dependable age-associated methylation marker, noting its resilience to environmental factors. Enhanced age estimation accuracy and reduced residual bias were observed in datasets incorporating the ELOVL2 marker. While the Dias et al. [12] dataset encompassed a broad age range of 1 to 93 years, the Zhou et al. [14] and Fakhr et al. [15] datasets included only individuals aged 20 and above. This discrepancy accounts for the linear relationship between age and ELOVL2 methylation observed in the Fakhr et al. dataset, contrasted with the non-linear relationship evident in the Dias et al. dataset (Fig. 2 & Fig. 4). Future research should prioritize model development that accounts for the age distribution characteristics reported in Dias et al., particularly within forensic applications. Such approaches are critical for the accurate identification of minors in migration contexts, where non-linear modeling techniques may yield more unbiased and precise age estimations.

ddPCR has emerged as a promising technique for forensic age estimation due to its robust performance with minimal DNA input and its capacity to deliver precise DNA methylation quantifications. The methodology developed herein constitutes a significant advancement in age prediction, offering improved accuracy and minimized bias, thereby serving as a comprehensive resource for forensic casework support. As the discipline progresses toward the practical application of these approaches, it is increasingly critical that such tools yield reliable and insightful data to aid investigative processes.

## Notes

### Competing Interest Statement

The authors have declared no competing interest.

