## Supplemental Figure 1 for "Improving the Accuracy of Forensic Age Estimation Through Bias Reduction"

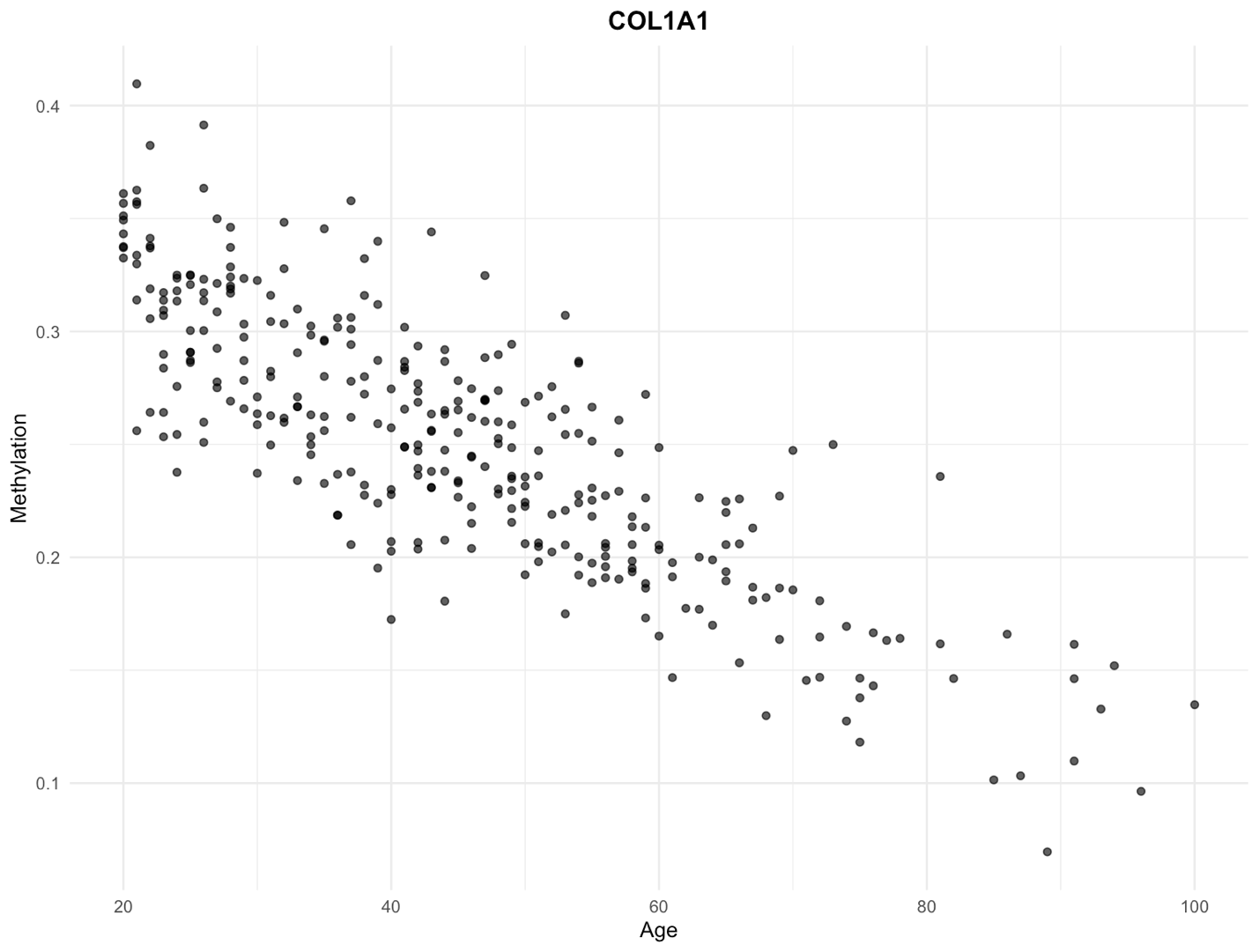

**Figure S1:** Scatterplot showing relationship between chronological age and methylation levels of marker COL1A1.
